# COVID-19 Associated Pulmonary Aspergillosis isolates are genomically diverse but similar to each other in their responses to infection-relevant stresses

**DOI:** 10.1101/2022.12.14.520265

**Authors:** Matthew E. Mead, Patrícia Alves de Castro, Jacob L. Steenwyk, Jean-Pierre Gangneux, Martin Hoenigl, Juergen Prattes, Riina Rautemaa-Richardson, Hélène Guegan, Caroline B. Moore, Cornelia Lass-Flörl, Florian Reizine, Clara Valero, Norman Van Rhijn, Mike J Bromley, Antonis Rokas, Gustavo H. Goldman, Sara Gago, the ECMM CAPA Study Group

## Abstract

Secondary infections caused by the pulmonary fungal pathogen *Aspergillus fumigatus* are a significant cause of mortality in patients with severe Coronavirus Disease 19 (COVID-19). Even though epithelial cell damage and aberrant cytokine responses have been linked with susceptibility to COVID-19 associated pulmonary aspergillosis (CAPA), little is known about the mechanisms underpinning co-pathogenicity. Here, we analysed the genomes of 11 *A. fumigatus* isolates from patients with CAPA in three centres from different European countries. CAPA isolates did not cluster based on geographic origin in a genome-scale phylogeny of representative *A. fumigatus* isolates. Phenotypically, CAPA isolates were more similar to the *A. fumigatus* A1160 reference strain than to the Af293 strain when grown in infection-relevant stresses; except for interactions with human immune cells wherein macrophage responses were similar to those induced by the Af293 reference strain. Collectively, our data indicates that CAPA isolates are genomically diverse but are more similar to each other in their responses to infection-relevant stresses. A larger number of isolates from CAPA patients should be studied to identify genetic drivers of co-pathogenicity in patients with COVID-19.

**Importance:** Coronavirus disease 2019 (COVID-19)-associated pulmonary aspergillosis (CAPA) has been globally reported as a life-threatening complication in some patients with severe COVID-19. Most of these infections are caused by the environmental mould *Aspergillus fumigatus* which ranks third in the fungal pathogen priority list of the WHO. However, little is known about the molecular epidemiology of *Aspergillus fumigatus* CAPA strains. Here, we analysed the genomes of 11 *A. fumigatus* isolates from patients with CAPA in three centres from different European countries and, carried out phenotypic analyses with a view to understand the pathophysiology of the disease. Our data indicates that *A. fumigatus* CAPA isolates are genomically diverse but are more similar to each other in their responses to infection-relevant stresses.

## Introduction

Lung coinfections and super infections caused by either bacteria or fungi are frequent and increase mortality in patients with severe COVID-19 (Gangneux *et al*. 2022; Hoenigl *et al*. 2022). Among fungal species known to cause secondary infections in patients already infected with severe acute respiratory syndrome coronavirus 2 (SARS-CoV2), *Aspergillus* species can give rise to COVID-19 Associated Pulmonary Aspergillosis (CAPA) in about 15.1% of ICU-admitted COVID-19 patients (Feys *et al*. 2021). However, the incidence of CAPA varies across medical centres and has been reported to range between 0.7 and 34.4%. Nevertheless, in a retrospective study of the literature (Salmanton-García *et al*. 2021), it was reported that 52.5% patients with CAPA died early after the diagnosis of the disease (<6 weeks after CAPA diagnosis) and 33.0% of these deaths were attributed to aspergillosis. Therefore, we need to improve our understanding of the molecular epidemiology of *Aspergillus fumigatus* CAPA strains with a view to better understand the disease and its impact on human health. (Steenwyk *et al*. 2021b).

Infections due to *A. fumigatus* are the most common cause of CAPA, but other *Aspergillus* species have been recently found in the clinic (Salmanton-García *et al*. 2021; Feys *et al*. 2021). Attempts to understand why *A. fumigatus* is the most frequent cause of aspergillosis have been carried out for decades. Several of these studies have focused on understanding traits related to its virulence in susceptible host (Nierman *et al*. 2005a; Fedorova *et al*. 2008; Abad *et al*. 2010; Brown and Goldman 2016; Bignell *et al*. 2016; Mead *et al*. 2021). Several studies have shown *A. fumigatus* phenotypic heterogeneity in infection-relevant traits and this has been linked to differences in virulence (Fuller *et al*. 2016; Kowalski *et al*. 2016; Ries *et al*. 2019). Moreover, phenotypic heterogeneity is largely attributed to genomic heterogeneity between *A. fumigatus* isolates (Barber *et al*. 2021; Rhodes *et al*. 2022; Horta *et al*. 2022) since approximately 16-42 % of the genome of an *A. fumigatus* isolate is variable (Barber *et al*. 2021; Lofgren *et al*. 2022).

Different mechanisms generate diversity and can facilitate adaptation to specific niche environments in *A. fumigatus*. For example, the generation of genetic variation in patients with chronic pulmonary infections has been linked to parasexual recombination (Engel *et al*. 2020) and the emergence of non-synonymous mutations (Ballard *et al*. 2018). Less is known regarding the heterogeneity of the phenotypes and genomes of *A. fumigatus* CAPA isolates as the disease is relatively new. To elucidate whether genomic- and pathogenicity-related characteristics in CAPA isolates are similar to non-CAPA, but clinically relevant isolates, we previously analyzed the genomic, chemical, and phenotypic heterogeneity of four CAPA isolates from Germany (Steenwyk *et al*. 2021b). Surprisingly, we found that the four CAPA isolates were more closely related to each other than to other *A. fumigatus* isolates and displayed only some degree of phenotypic heterogeneity. Aiming to better understand whether this lack of genomic diversity holds true across CAPA isolates, we built upon our previous study and performed genomic and phenotypic traits analyses of 11 additional *A. fumigatus* CAPA isolates from three European centres (Graz, Manchester, and Rennes). We observed that *A. fumigatus* CAPA isolates are genomically diverse but are more similar to each other in their responses to infection-relevant stresses. We conclude that *A. fumigatus* CAPA isolates likely span the genomic and phenotypic diversity of *A. fumigatus*.

## Materials and Methods

### Patient information and ethics approval

*Aspergillus fumigatus* isolates were obtained as part of the multinational CAPA observational study of the European Confederation of Medical Mycology (Prattes *et al*. 2022). Each participating study centre (Manchester, Graz, and Rennes) was responsible for obtaining local institutional review board and/or local ethics policy approval. Institutional review board approval numbers are as follows: Medical University of Graz EC #32-296 ex 19/20; at the University of Manchester data acquisition was conducted as a retrospective audit, which does not require local ethics but was approved by the hospital’s audit committee; at Rennes University Hospital this protocol was approved by the local ethics committee (approval number, 20.56). The study has been performed in accordance with the ethical standards laid down in the 1964 Declaration of Helsinki and its later amendments.

### DNA extraction and sequencing

All 11 *A. fumigatus* CAPA isolates **(Table 1)** were grown from 1×10^7^ asexual spores (conidia) in minimal media (MM) [1% (w/v) glucose, nitrate salts, trace elements, pH 6.5] ((Barratt *et al*. 1965)Käfer, 1977) for 20 hrs at 37°C. After mechanical disruption of the mycelia, genomic DNA extraction was performed in phenol: chloroform (1:1). DNA quantity and quality were assessed using a NanoDrop 2000 spectrophotometer (Thermo Scientific). The DNA purity ranged from 1.8 to 2.0 for OD260/280 and 2.0-2.2 for OD260/230.

**Table 1.** Patient and isolate information.

| Isolate Name | City/Country of Origin | Underlying Condition | Age | Gender | Sample Origin |
| --- | --- | --- | --- | --- | --- |
| CAPA 1 | Rennes, France | Chronic myeloid leukemia, ARDS | 79 | Female | BAL + Tracheal aspiration |
| CAPA 2 | Rennes, France | Chronic myelo-monocytic leukemia, ARDS | 78 | Female | Tracheal aspiration |
| CAPA 3 | Rennes, France | ARDS | 75 | Male | Tracheal aspiration |
| CAPA 4 | Rennes, France | ARDS | 58 | Male | Tracheal aspiration |
| CAPA 5 | Rennes, France | ARDS | 78 | Female | Tracheal aspiration |
| CAPA 6 | Rennes, France | ARDS, Obesity, Hypertension | 71 | Male | Tracheal aspiration |
| CAPA 7 | Rennes, France | ARDS, Hypertension | 73 | Male | Tracheal aspiration |
| CAPA 8 | Graz, Austria | ARDS | 65 | Male | Tracheal aspiration |
| CAPA 9 | Graz, Austria | ARDS | 60 | Male | BAL |
| CAPA 10 | Manchester, UK | ECMO | 41 | Male | BAL |
| CAPA 11 | Manchester, UK | ECMO | 41 | Male | BAL |

ARDS – Acute Respiratory Distress Syndrome, CAPA – COVID-19 associated pulmonary aspergillosis, ECMO – Extra-Corporeal Membrane Oxygenation. BAL: bronchoalveolar lavage. Note that CAPA 10 and 11 were isolated from the same patient.

Library preparation and sequencing was carried out by Vanderbilt Technologies for Advanced Genomics (VANTAGE). Libraries were prepared using the NEBNext Ultra II DNA Library Prep Kit. Sequencing of the libraries was carried out on an Illumina NovaSeq 6000 to produce paired-end, 150bp reads.

### *De novo* genome assembly, annotation, and quality determination

To obtain high-quality and adapter-free reads, raw reads were trimmed with Trimmomatic version 0.39 (Bolger *et al*. 2014) using the parameters “2:30:10 LEADING:3 TRAILING:3 SLIDINGWINDOW:4:15 MINLEN:36”. On average, 36 million read pairs passed trimming. Trimmed reads were then assembled with SPAdes version 3.15.2 (Prjibelski *et al*. 2020) using the parameters “--isolate” and “--cov-cutoff auto”. Genome statistics were calculated with BioKIT version 0.0.4 (Steenwyk *et al*. 2022).

To identify putative protein-coding genes, Augustus version 3.3.2 (Stanke *et al*. 2008a) was used to annotate the newly assembled CAPA genomes. The *Aspergillus fumigatus* annotation that is packaged with the software was used as a training dataset. Completeness and fragmentation of the genomes was determined with version 4.0.4 of BUSCO (Manni *et al*. 2021) using the default Eurotiales database. All quality metrics for the genome assemblies and annotations of the new CAPA isolates were comparable to values for reference strains Af293 and A1163 (Nierman *et al*. 2005b; Fedorova *et al*. 2008; Amos *et al*. 2022).

### Phylogenomic tree inference

To determine the general taxonomy of the 11 new CAPA isolates, we built a single gene tree of the *tef1* homologs from the new CAPA strains and the 100 genes most-similar to the Tef1 ortholog present in *A. fumigatus* strain Af293 (<u>XM_745295.2</u>). Homologs of *A. fumigatus* Tef1 were identified in both the CAPA isolates and the NCBI nucleotide collection (nr/nt database) with blastn verion 2.8.1(Altschul *et al*. 1990) using default parameters. The 112 *tef1* sequences (11 CAPA + 100 NCBI + 1 *A. fumigatus* Af293) were aligned with MAFFT version 7.402 (Katoh *et al*. 2002; Katoh and Standley 2013) and the following parameters: -op 1.0 -maxiterate 1000 -retree 1 -genafpair. The resulting alignment was trimmed with version 1.2.0 of ClipKIT (Steenwyk *et al*. 2020) and a tree was made from the trimmed alignment using version 1.6.12 of IQ-TREE with the model finder parameter and 5,000 ultrafast bootstraps (Nguyen *et al*. 2015a). A cladogram of the tree was visualized with iTOL version 5 (Letunic and Bork 2021a).

To infer the phylogenetic relationships between the 11 new CAPA isolates and other *A. fumigatus* strains, a modified version of a previously published pipeline was employed (Steenwyk *et al*. 2021b). Specifically, genomes from 50 taxa (three non-*A. fumigatus* outgroup strains, 43 *A. fumigatus* isolates that span the diversity of the species, and four previously analyzed CAPA isolates) were obtained from our previous study (https://doi.org/10.6084/m9.figshare.13118549 (Steenwyk *et al*. 2021b)) and compared to the genomes of the 11 new CAPA isolates.

To discover suitable loci for phylogenetic reconstruction, 4,525 single-copy orthologs identified amongst the 50 previously analyzed genomes were obtained. A Hidden Markov Model (HMM) was made for each orthologous group of genes using hmmbuild within HMMER version 3.2.1 (hmmer.org). These 4,525 HMMs were used as input in orthofisher version 1.0.3 (Steenwyk and Rokas 2021) to identify the copies of the orthologs in the genomes of the 11 new CAPA isolates with the parameter “-b 0.95”. Ten of the orthologs were found to vary in their copy number across the 11 new genomes and were not used in subsequent analyses. Protein sequences from the 4,515 single-copy orthologs from all 61 taxa were combined into 4,515 FASTA files for further analysis.

To align the 4,515 single-copy orthologs, MAFFT version 7.402 (Katoh *et al*. 2002; Katoh and Standley 2013) was used along with the parameters “-bl 62 -op 1.0 -maxiterate 1000 -retree 1 - genafpair” (Steenwyk *et al*. 2022). The 4,515 alignments were trimmed with version 1.2.0 of ClipKIT (Steenwyk *et al*. 2020) and then combined into a supermatrix with PhyKIT version 1.5.0 (Steenwyk *et al*. 2021a). The resulting supermatrix contained 2,361,569 amino acid sites and was analyzed using IQ-TREE version 1.6.12 (Jones *et al*. 1992; Nguyen *et al*. 2015b) and parameters “-bb 5000 -m TEST -nbest 10 –runs 5 -safe” to produce a maximum likelihood tree. Note that these parameters included using 5,000 ultrafast bootstrap support approximations (Hoang *et al*. 2018). The tree was visualized with iTOL version 5 (Letunic and Bork 2021b).

### Antifungal susceptibility testing

Antifungal susceptibility testing of the CAPA isolates was performed using the EUCAST (European Committee for Antimicrobial Susceptibility Testing) reference microdilution method version 9.3.2 (Arendrup *et al*. 2020) For all isolates, susceptibility to amphotericin B, isavuconazole, voriconazole, posaconazole, amphotericin B, caspofungin and micafungin was tested at 48 hrs.

### Growth assays

*Aspergillus fumigatus* radial growth from CAPA isolates and the reference strains Af293 and A1160 was comparatively analyzed on either solid Minimal Media (MM) or MM supplemented with different concentrations of stressor agents (sorbitol (1M), Congo Red (10 mg/ml), or hydrogen peroxide (1.5 mM)) at 37 °C. For the different iron availability conditions, iron was omitted from the trace element solution (Käfer 1977) and supplemented at various concentrations (3 µM for iron depletion or 300 µM for iron excess). The ferreous iron chelator bathophenanthroline disulfonic acid (BPS) was used at 200 µM to increase iron starvation in solid media as described in (Gsaller *et al*. 2014). Plates were inoculated with 10^4^ spores per strain and growth was then measured after 72 hrs. Radial growths were expressed as ratios, dividing colony radial diameter of growth in the stress condition by colony radial diameter in the control (no stress) condition. The capacity to grow under hypoxia (5% CO_2_ 1% O_2_) and at 44 °C was also evaluated. Experiments were done using two or more biological and technical replicates. Statistical comparisons of growth rate of the CAPA isolates versus reference strains Af293 and A1160 were done using two-way ANOVA with Dunnett’s post-hoc test (GraphPad Prism v9, La Jolla, CA).

### Pathogenicity assays

To investigate differences in *A. fumigatus* killing by macrophages, 10^6^ RAW 264.7 cells were seeded in 6-well plates and incubated for 24 hrs. RAW 264.7 murine macrophages (ATCC TIB-71) were maintained at 37 °C, 5% CO_2_ in Dulbecco’s Modified Eagle’s Medium (DMEM) supplemented with 10% fetal bovine serum (FBS) and a 1% Penicillin – Streptomycin solution, all from Merck (Darmstadt, Germany). Macrophages were used under passage 20. Cells were then challenged with 10^6^ spores of each of the isolates and incubated for 6 hrs. Cells were then lysed in water and plated in Sabouraud Agar Plates. Colony Forming Units (CFUs) were enumerated after 24 hrs of incubation at 37°C. To correct for strain heterogenicity, the number of CFUs for a particular isolate in confrontation experiments with macrophages was divided by the number of CFUs for that isolate in the absence of macrophages. Experiments were done using three or more biological replicates and technical duplicates. Statistical comparisons of macrophage killing of *A. fumigatus* CAPA isolates and the reference strains A1160 and Af293 were done by One-Way ANOVA with Dunnett’s post-hoc tests (GraphPad Prism v9, La Jolla, CA).

It has been previously described that *A. fumigatus* germination is critical to induce cytokine responses and cytotoxicity of host cells during infection. To investigate whether *A. fumigatus* CAPA isolates were able to induce host cell damage and activate macrophage responses in a different manner than the reference strains Af293 and A1160, 10^6^ RAW 264.7 macrophages were seeded in 24 well-plates and challenged with 10^6^ spores (Furukawa *et al*. 2020) for 9 and 24 hrs. Lactate dehydrogenase (LDH) release was measured using the Cyto Tox 96 © Non-Radioactive Cytotoxicity Assay (Promega, Madison, Wisconsin, USA) according to manufacturer’s instructions. The concentrations of IL-6 and TNF alpha in cell culture supernatants was measured by using the Mouse IL-6 and TNF alpha DuoSet ELISA (R&D systems, Minneapolis, Minnesota, USA). Statistical differences in LDH release and cytokine secretion between RAW 264.7 macrophages challenged with *A. fumigatus* CAPA isolates and reference strains were determined by one-way multiparametric ANOVA with Dunnett’s correction using GraphPad Prism 9.0 (La Jolla, CA, USA).

### Data Availability

Assembled genomes and annotations used in this study are available via FigShare at https://figshare.com/articles/dataset/COVID-19_Associated_Pulmonary_Aspergillosis_Isolates_are_genomically_diverse_but_are_more_similar_to_each_other_in_their_responses_to_infection-relevant_stresses/20409096. Reads, assemblies, and annotations that met NCBI formatting guidelines and are very similar to those discussed here, are available through BioProject PRJNA787571. Note that for the NCBI genomes “Sample #” is synonymous to “CAPA #”.

## Results and Discussion

### New CAPA isolates represent diverse lineages of the *A. fumigatus* phylogeny

To determine the evolutionary relationships between the 11 newly identified and sequenced CAPA isolates, four previously analyzed CAPA isolates, 55 *A. fumigatus* strains, and three outgroup taxa (two strains of *A. fischeri* and one of *A. oerlinghausenensis*, the closest known relatives of *A. fumigatus* (Houbraken *et al*. 2016; Steenwyk *et al*. 2020b)), were used to infer the phylogeny of these strains **(Figure 1, Supplementary Table 1 10.6084/m9.figshare.21688172)**. Our tree showed that the 11 new CAPA isolates belonged to *A. fumigatus*; however, these 11 new isolates were more diverse than the four previously sequenced CAPA isolates. The four previously sequenced CAPA isolates all originated from Germany and were very closely related to each other and to the strains A1163 and Af293 (Steenwyk *et al*. 2021b). In contrast, none of the new CAPA isolates are closely related to A1163 and Af293 or the previously sequenced CAPA isolates; instead, these isolates span the *A. fumigatus* phylogeny.

**Figure 1.**
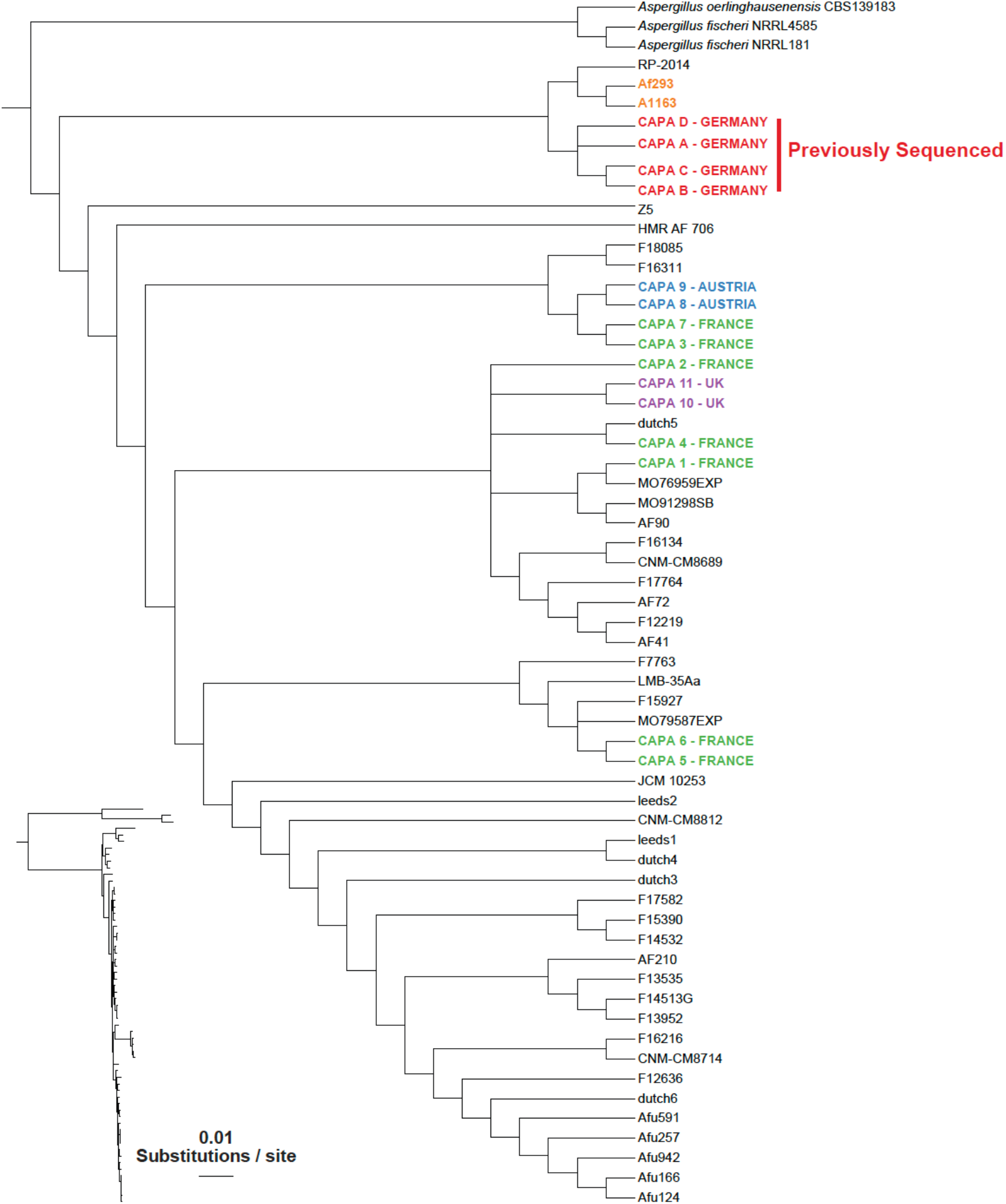
Phylogeny of 55 *A. fumigatus* isolates and 3 outgroup taxa reveals that the 11 new CAPA isolates span the genomic diversity of the species. 4,515 single-copy orthologs from a total of 61 taxa (11 new CAPA isolates, three non-*A. fumigatus* outgroup strains, 43 *A. fumigatus* isolates that span the diversity of the species, and four previously analyzed CAPA isolates) were used as input to construct a maximum likelihood tree. A slight geographic clustering of CAPA isolates was observed but isolates from different locales were more diverse than a previous set of four CAPA isolates (all of which were isolated from the same hospital in Germany).

Two *A. fumigatus* CAPA isolates from Austria and two CAPA isolates from a patient from the U.K. were most closely related to each other, respectively. However, the CAPA isolates from France exhibited more genomic diversity and were located across a greater portion of the phylogeny. Our previously sequenced CAPA isolates (A, B, C, and D) and newly sequenced isolates CAPA 1, 2, and 4, were sister with non-CAPA *A. fumigatus* isolates, suggesting that there are likely to be a few or no genomic traits that are uniquely shared only by CAPA isolates. Interestingly, this finding is in disagreement with our previously published data showing that CAPA isolates from the same geographic area are closely related, thus suggesting a possible common source of infection (Steenwyk *et al*. 2021b).

### Strain heterogenicity of CAPA isolates in virulence-related culture conditions and antifungal drug susceptibility

*A. fumigatus* isolates from CAPA patients displayed strain-dependent variation in growth phenotypes when compared to the reference strains Af293 and A1160 **(Figure 2 and Supp Figure 1, 10.6084/m9.figshare.21688172)**. In general, most CAPA isolates displayed phenotypes similar to the reference strain A1160 when grown under hypoxia, osmotic stress, high temperature (44 °C) or low and high concentrations of iron. For three of the CAPA isolates, radial growth in the presence of cell wall stress (CAPA 6), oxidative stress (CAPA 7 and 9), and/or iron starvation stress (CAPA 6 and 7) was significantly reduced compared to both Af293 and A1160 (two-way ANOVA with Dunnett’s post-hoc test, P < 0.05). Reduced sensitivity to cell wall damaging agents was not detected in the 11 CAPA isolates included in this study. However, we used Congo Red as a stressor rather than calcofluor white, which was used in (Steenwyk *et al*. 2021b). All CAPA isolates grow similarly to *A. fumigatus* reference strains when cultured in solid MM (without any stress). All isolates were susceptible to all antifungal drugs included in the study; posaconazole displayed minimal inhibitory concentrations close to resistance breakpoints defined, but when retested never exceeded MICs > 0.25 **(Table 2)**.

**Figure 2:**
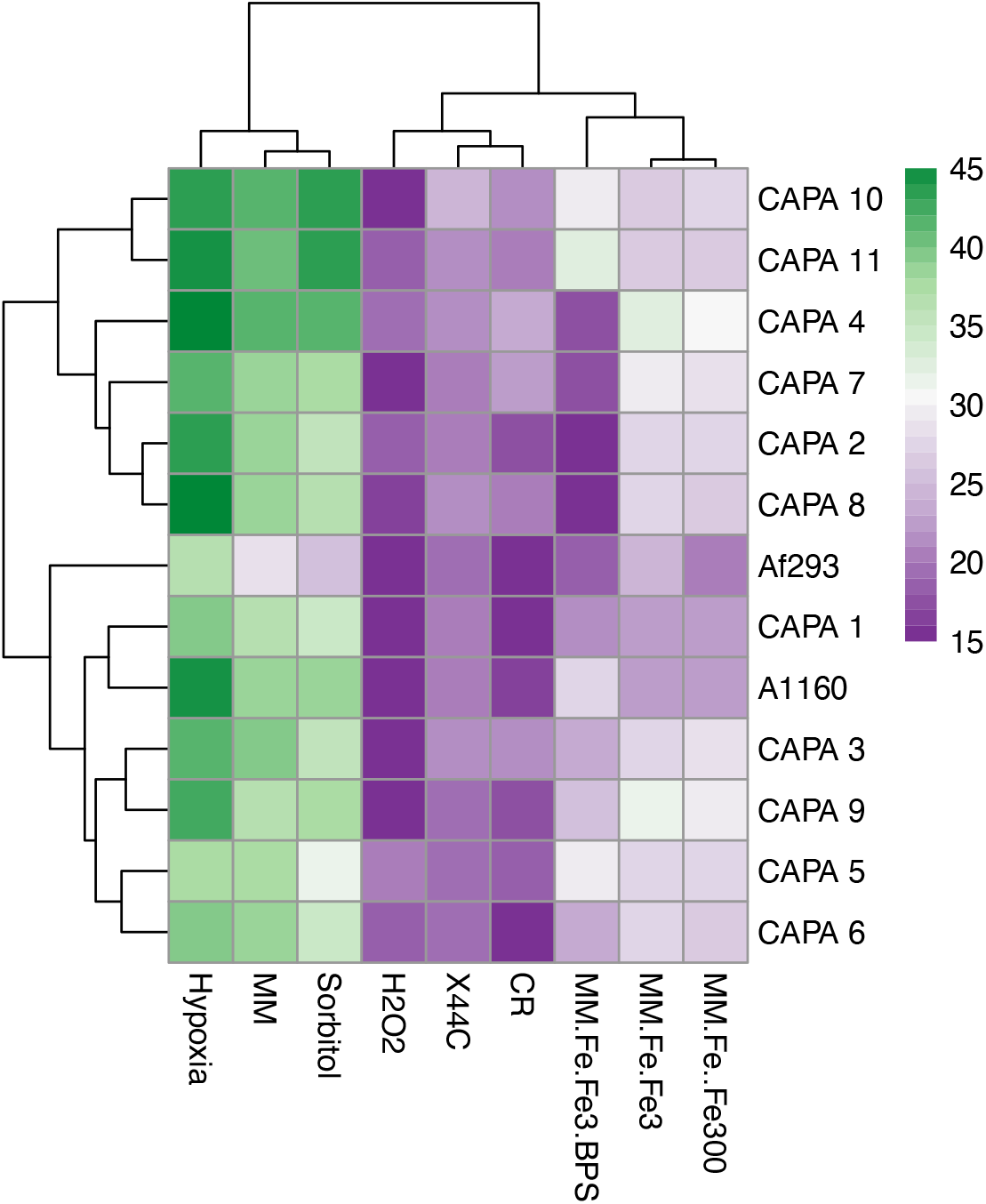
*A. fumigatus* radial growth in infection-relevant culture media. Data is represented as mean of colony diameter (mm) of *A. fumigatus* CAPA strains and controls. Clustering of strains was carried out according to diameter size.

**Table 2.** Susceptibility profile of the 11 CAPA isolates included in this study at 48 h. AMB: Amphotericin B, CSP: caspofungin, MCF: micafungin, PCZ: Posaconazole, VCZ: voriconazole, IVZ: isavuconazole. Isolates were tested twice and duplicates. MICs given were converted to the highest concentration detected.

| Isolate | MIC (mg/L) |  |  |  |  |  |
| --- | --- | --- | --- | --- | --- | --- |
|  | AMB | CSP | MCF | PCZ | VCZ | IVZ |
| **1** | 0.5 | 0.125 | 0.032 | 0.25 | 1 | 0.5 |
| **2** | 0.5 | 0.125 | 0.032 | 0.25 | 1 | 1 |
| **3** | 0.5 | 0.125 | 0.032 | 0.25 | 1 | 1 |
| **4** | 0.5 | 0.125 | 0.032 | 0.25 | 0.5 | 1 |
| **5** | 0.5 | 0.125 | 0.032 | 0.25 | 1 | 1 |
| **6** | 1 | 0.125 | 0.032 | 0.25 | 1 | 1 |
| **7** | 1 | 0.25 | 0.032 | 0.25 | 0.5 | 0.5 |
| **8** | 0.5 | 0.25 | 0.032 | 0.25 | 1 | 1 |
| **9** | 0.5 | 0.25 | 0.032 | 0.25 | 1 | 1 |
| **10** | 0.5 | 0.25 | 0.032 | 0.25 | 1 | 1 |
| **11** | 0.5 | 0.25 | 0.032 | 0.25 | 1 | 1 |

### CAPA isolates are more efficiently killed by macrophages than reference strain A1160

Macrophage killing of *A. fumigatus* conidia is one of the main mechanisms of antifungal defence during infection. Pulmonary macrophages in COVID-19 have been described to be hyperactivated thus favouring tissue damage at the site of infection (Knoll *et al*. 2021). The efficiency of macrophages to kill *A. fumigatus* CAPA isolates and two other reference isolates was comparatively analyzed at 6 hours post-infection. CAPA isolates were less susceptible to macrophage killing compared to reference strain A1160 but exhibited killing rates on par to those of reference strain Af293 (**Figure 3a**, P< 0.05). *In vivo* studies using immunosuppressed mouse models of infection and neutrophil-depleted zebra fish larvae have shown that *A. fumigatus* strain CEA10 is more virulent than Af293 (Kowalski *et al*. 2016; Knox *et al*. 2016; Caffrey-Carr *et al*. 2017). However, our data indicate decreased killing of *A. fumigatus* Af293 *in vitro*. Moreover, it has been recently reported that *A. fumigatus* Af293 might be more pathogenic in immunocompetent hosts (Rosowski *et al*. 2018). Differences in *A. fumigatus* killing were not correlated with differential cytokine profiles at 9 h post infection **(Figure 3b-c)**. Only *A. fumigatus* CAPA isolates 2 and 10 showed increased secretion of IL-6 and/or TNF alpha compared to any of the control strains at 9 h post infection. Differences in the capacity of macrophages to kill CAPA isolates compared to *A. fumigatus* controls was not linked to decreased cell cytotoxicity as measured by LDH release **(Figure 3d)**. Altogether, these data indicate that the CAPA isolates included in this study have generally similar pathogenicity profiles during infection with macrophages to the reference strain Af293.

**Figure 3.**
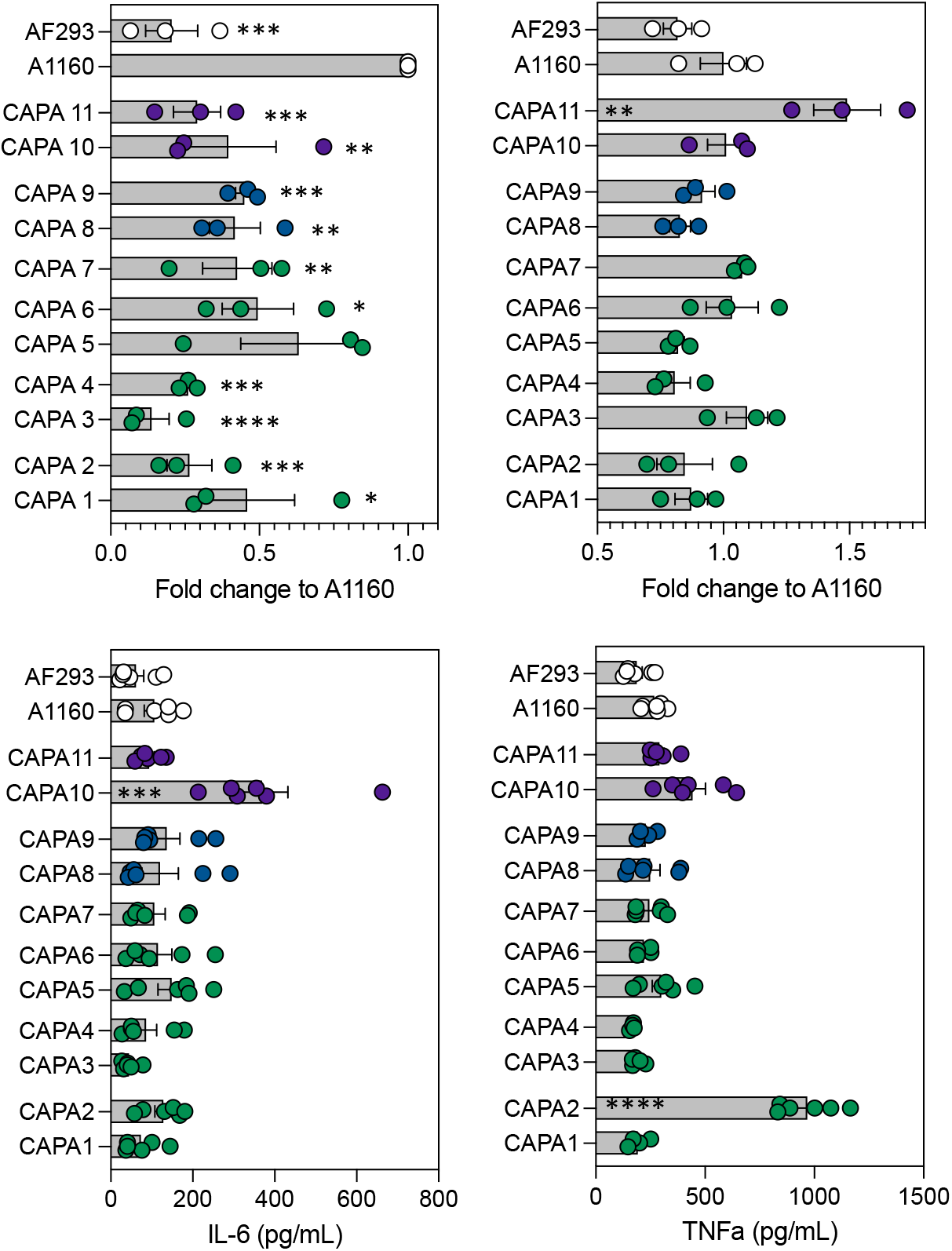
CAPA isolates exhibit in vitro RAW 263.7 macrophages responses similar to Af293. **(a)** Fold change killing of *A. fumigatus* CAPA isolates and reference strains Af293 and A1160 by RAW 263.7 macrophages at 6 h post infection. **(b)** LDH release of macrophages challenged with 11 CAPA isolates and reference strains Af293 and A1160 at 24 h post infection. (c) IL-6 (d) TNFa release by RAW 263.7 macrophages at 9 h post-infection with CAPA strains and controls. Data represent mean and standard deviation of a minimum of three biological and technical replicates. Neat controls were subtracted from test samples and used for background corrections. * P < 0.05, ** P < 0.01, ***P<0.001 and ***P<0.0001 compared to A1160.

In this study, we found *A. fumigatus* isolates from patients with CAPA are genetically heterogenous but phenotypically similar. Increasing the number of available CAPA genomes since our previous study has allowed us to observe that CAPA isolates represent diverse lineages of the *A. fumigatus* phylogeny and the concomitant absence of geographical clusters, which contrasts with our earlier findings based on an analysis of four CAPA isolates from Germany (Steenwyk *et al*. 2021b). The 11 new CAPA isolates have similar phenotypic profiles to *A. fumigatus* A1160 when tested for sensitivity to stressors in culture-relevant conditions, while macrophage resistance phenotypes were more similar to Af293. However, this may be an artefact due to the relative small number of isolates included in the study (our previous study indicated that CAPA isolate secondary metabolite profiles were more similar to *A. fumigatus* Af293 than to A1160 (Steenwyk *et al*. 2021b). Recent work suggests that differences in virulence for *A. fumigatus* A1160 and Af293 might be determined by the experimental system used (Bertuzzi *et al*. 2021). However, recent data suggest that *A. fumigatus* Af293 but not A1160 triggers SARS-CoV2 replication within airway epithelial cells (Dancer *et al*. 2022); however, what specific fungal factors are critical regulators of co-pathogenesis in CAPA remains to be determined. In-host microevolution of *A. fumigatus* strains during chronic infections has been reported in the literature (Ballard *et al*. 2018; Engel *et al*. 2020). Even though this has only been reported in long-term infections, a previous study attempted to determine whether specific SNPs or copy number variants in genetic determinants of virulence and biosynthetic gene clusters could explain *A. fumigatus* CAPA genomic heterogeneity (Steenwyk *et al*. 2021b). An early stop codon in *pptA* was found, but this did not correlate with reduced production of secondary metabolites in CAPA isolates. Continued genome sequencing of additional CAPA isolates, such as those we report here, may facilitate identifying mutations that impact infection-relevant traits.

## FUNDING/ACKNOWLEDGENMENTS

LS and AR were funded by the Howard Hughes Medical Institute through the James H. Gilliam Fellowships for Advanced Study program. Research in AR’s lab is supported by grants from the National Science Foundation (DEB-2110404), the National Institutes of Health/National Institute of Allergy and Infectious Diseases (R56 AI146096 and R01 AI153356), and the Burroughs Wellcome Fund. SG was co-funded by the NIHR Manchester Biomedical Research Centre, The Fungal Infection Trust, Manchester Academy of Health Sciences and The Dowager Countess

Eleanor Peel Trust. MB and NvR were supported by the Wellcome Trust grant number 219551/Z/19/. We thank the Fundação de Amparo à Pesquisa do Estado de São Paulo (FAPESP) grant number 2021/04977-5 (GHG) and BEPE 2020/01131-5 (CV) and the Conselho Nacional de Desenvolvimento Científico e Tecnológico (CNPq) grant numbers 301058/2019-9 and 404735/2018-5 (G.H.G.) and 163550/2020-4 (PAC), both from Brazil, and the National Institutes of Health/National Institute of Allergy and Infectious Diseases grant R01AI153356 (GHG).

## Supplementary Material (available on 10.6084/m9.figshare.21688172)

**Supplementary Figure 1. Radial growth ratios of the 11 CAPA isolates and reference strains Af293 and A1160 under hypoxia, at 44 °C, and in the presence of osmotic, cell wall, oxidative, and iron stress conditions**. Growth differences between CAPA isolates and reference strains Af293 (black) and A1160 (red) were analysed by using two-way ANOVA with multiple comparison test. * P < 0.05.

**Table S1. Genomic information for the 11 *A. fumigatus* CAPA isolates sequenced in this study**.

